# Microembossing Hydrogel Meso-Circuits for Patterning Dissociated Neurons Promotes Ensemble Formation*

**DOI:** 10.64898/2026.03.03.709093

**Authors:** Mckennah Thompson, Connor L. Beck, Anja Kunze

## Abstract

Functional networks of wired neurons comprise the basis for neuronal computation and processing. Within neuronal networks, activation of unique ensembles is an important identity of neuronal processing. However, dissociated neuronal networks form homogeneous functional structures with minimal variety in ensemble dynamics. To reintroduce such dynamics, we propose structuring the networks to follow multi-connectivity (micro- and meso-network) paradigms. Here, we use agarose microembossing to physically pattern dissociated neuronal networks across these scales. To perform agarose microembossing, we impress features with poly-dimethyl-siloxane (PDMS) stamps into liquid agarose to emboss features which hold under cold gelation. We validate the viability of primary neurons within the hydrogel patterns and interrogate circuit dynamics through calcium imaging. Patterned features presented with robust ensemble dynamics that are dependent on connectivity paradigms. Altogether, this work establishes a platform for investigating how engaging multi-scale features in the physical network informs neuronal ensemble dynamics.

**Clinical Relevance:** This work enables further dissociated studies to probe dynamics. We expect that this platform would be especially useful in early-stage drug development or personalized medicine pipelines that need to investigate circuit dynamics.

## I. Introduction

Understanding the intricate interplay of neuronal networks within the cerebral cortex is a fundamental pursuit of neuroscience. These networks are comprised of neurons that form connectivity across scales, establishing the foundation for a multitude of computations that underlie the decision-making processes in mammals. Neurons establish connections at multiple scales, from local connectivity of microcircuits to intermediate meso-circuit connectivity patterns that interact with global macro-circuits to perform complex tasks [1]. How such connectivity paradigms inform function in neuronal networks remains a critical barrier. To elucidate such interactions, isolating independent neuronal networks can provide key insights. However, functional isolation can be particularly difficult *in vivo*, given the pervasive inter-connectivity of the established networks in the brain. To circumvent this, dissociated *in vitro* networks allow for the complete isolation of neurons by enzymatic digestion and trituration. These isolated neurons, seeded into culture environments, regrow neurites that will mature into axons and dendrites, driving neuronal polarity and establishing functional connections with neighboring cells.

As neurons form functional connections in vitro, the random seeding of cellular position and uninformed growth cues leads to haphazard neurite outgrowth, resulting in a randomly formed network. It has been shown that these random networks can produce rich-club connectivity like the brain [2], suggesting that dissociated neurons have the potential to configure in brain-like states. However, there is still a large separation between the applicability of dissociated circuits to that of their *in vivo* counterparts. We expect this divergence in dissociated cultures is likely due to the lack of scaled connectivity [3], which is formed during brain development. Specifically, *in vivo* cortical circuits process information through ensembles, which are local clusters of neurons with high connectivity, that can process information propagated by global circuits [4]. Therefore, by introducing brain-like connectivity scales into dissociated cultures, we hypothesize that the structuring of neuronal networks will generate functional patterns more closely related to the brain.

To engineer dissociated circuits, various tools have been developed for improved consistency and reproducibility. Primarily, tools such as topographical or chemical cues have enabled engineering of directional neuronal circuits [5]. Traditionally, narrow microfluidic channels are used to guide axonal growth, preventing neuronal cell bodies from migrating between chambers [6]. These techniques have proven effective in creating directional cultures for engineering basic circuitry but are constrained by microchannel delivery. Using similar techniques, researchers have achieved resolutions ranging from small populations to single neurons [7], [8]. Alternatively, cell adhesion molecules, known for promoting neuronal growth, have been patterned using microfluidic techniques [9] and microcontact printing [10]. More recently, microembossing surface features in hydrogels has emerged as a method to topographically pattern neural circuits [11], [12]. Engineering dissociated cortical circuits has advanced our understanding of fundamental neurocomputation principles by limiting structural connectivity into linear components through axonal isolation. However, growing evidence suggests that the linear structuring of *in vitro* circuits lacks a crucial component of native cortical features. Thus, it has been postulated that defining multiple scales of connectivity within a dissociated network could enable function resembling features within the brain [3]. This paradigm has led to the idea of integrating multiple scales on a single chip, presenting the feasibility of such devices [13]. Consequently, the next generation of brain-on-chip technologies must preserve this multiscale structure.

In this work, we develop and characterize agarose embossing to structure dissociated neuronal networks with multi-scaled local- and meso-connectivity. We introduce robust patterning techniques which can be implemented across labs and perform validation of neuronal growth in the device. Finally, we use calcium imaging to explore how the networks behave functionally under spontaneous activity.

## II. Methods

### A. Agarose meso-circuit fabrication

To employ patterned neural circuits *in vitro*, we used hydrogel embossing through poly-dimethly-siloxane (PDMS) stamping following our previous work [12]. PDMS stamps were fabricated from silicon wafers, following a previous cleanroom fabrication protocol. In brief, a negative pattern photolithography mask was designed using CleWin (PhoeniX, Netherlands). KMPR 1025 (Microchem) was spin-coated (500 rpm for 5 s ramp-up speed with 100 rpm s^-1^ acceleration; 2000 rpm for 30 s; 500 rpm for 20 s ramp down speed) and soft-baked at 100 °C for 15 min to obtain a feature height of 50 μm. The features were exposed (700 mJ cm^−3^) to ultra-violet light using a contact aligner (Shipley SPR 1813) and developed (SU-8 developer) for 5 min followed by a thorough rinse with deionized (DI) water and triple solvent (acetone, isopropanol, ethanol).

The agarose meso-circuit fabrication began by designing and fabricating poly-dimethyl siloxane (PDMS) stamps. PDMS stamps were cast from micropatterned silicon wafers. 30g silicone elastomer base and 3g curing agent (Sylgard) were mixed using a 10:1 (w/w) base/curing agent ratio and poured onto the patterned wafer to cast the PDMS stamp. The PDMS elastomer mixture was cured for 2 h at 65 °C and gently peeled off from the master and cut to size. A 1% agarose in phosphate buffered saline (PBS) solution was prepared and added to a falcon tube 5x greater than the volume and vortexed, creating a cloudy white solution. PDMS stamps were placed in the autoclave with agarose simultaneously at 121 ºC and 1 bar for 15 min, with 15 min ramp times. Following autoclaving, the stamps were relocated to the biosafety cabinet (BSC) and agarose was maintained at 80 ºC in a dry oven. Magnetic clamps were sprayed with ethanol and added to the BSC along with 35 mm petri dishes. PDMS features were dropwise coated in poly-d-lysine (PDL, 0.5 ug/mL), with approximately 1 mL per stamp. A petri dish was placed in the magnetic clamp baseplate and agarose (80 ºC, 250 µL) was added to the dish. The stamp was added on top with the features facing down and the top of the magnetic clamp was carefully added to compress the stamp into the agarose with the magnetic force performing the work. The assembly was then placed in the refrigerator at 8 ºC for the specified time (14 to 32 min). After cooling, the apparatus was relocated to the BSC, the magnetic clamp was removed, and the stamp was carefully lifted off with tweezers. Devices were stored with PBS or culture media and kept in the incubator until seeding. In cases where fluorescent microparticles (1 μm, polystyrene, glacial blue, 1% solids) were used, microparticles were added following the agarose autoclaving (10 μL/mL) vortexed thoroughly, and heated at 80 ºC for 20 min.

### B. Cell culturing

We cultured dissociated primary cortical neurons from rat embryonic brain tissues (E18, TransnetYX). Neuronal cell cultures were established by following previously reported protocols by the authors. In brief, cortical hemispheres were dissected from whole tissues in phosphate buffered solution (PBS) with 33 mM glucose (1% (v/v) penicillin-streptomycin) and dissociated using 10% (v/v) papain (Carica papaya, Roche, pH 7.3, 15 min, 37 °C). Following dissociation, the papain solution was removed, a transfer solution (90% Neurobasal Plus, 10% Horse Serum, 37 ºC, Gibco) was added, and the cells were mechanically separated by triturating through a 1000 µL pipet. The dissociated cells were centrifuged (6 min, 500 rpm, at room temperature) to form a cell pellet. The supernatant was removed, and the cell pellet was resuspended in 2 mL of culture media (97% Neurobasal Plus, 2% B-27 Plus, 1% Glutamax, v/v, 37 ºC, Gibco). Neuronal cells were seeded at a concentration of 106 cells into 2 mL of culture media and incubated (37 °C, 95% air, 5% CO2, Relative Humidity), with culture media supplemented with penicillin-streptomyosin (96% Neurobasal Plus, 2% B-27 Plus, 1% PenStrep, 1% Glutamax, v/v, 37 ºC, Gibco) starting on 2 DIV and exchanged every two to three days.

### C. Live-dead staining

To identify viable neurons in agarose patterned microcircuits, cells were cultured to 8 DIV and a live/dead assay (R37601, Thermo) was used per manufacturers specification. The vials were thawed at 37 ºC in dry bath and 1 mL of the live (calcein AM) solution was added to the dead (BOBO-3 Iodide) lyophilized stock vial. The vial was sealed, vortexed for 30 s, and added to the neuronal cultures at 100 µL per Petri Dish. Each dish was incubated for at least 20 min before imaging. Cells were imaged with brightfield microscopy and fluorescently imaged for live (Calcein AM, GFP 488/520 nm ex/em) and dead (BOBO-3 Iodide, TXR 570/600 nm ex/em) cells. Post-experiment, Live/Dead count was analyzed in ImageJ. The brightfield image was used to determine the location of microembossed wells and the surface of agarose. Cells were manually counted in the green channel and red channel to determine the number of viable cells.

### D. Calcium imaging and ensemble detection

To record neuronal dynamics, we employed epifluorescent imaging with chemical calcium indicators. Cortical neurons (14 DIV) were exposed to a mixture of Fluo-4 AM (1:1 v/v ratio, Gibco) and Calbryte 520AM (5 μM) for at least 1 h per manufacturer specification. Videos were recorded with an inverted fluorescent microscope (Leica DMI-8, GFP 488nm, 200x total magnification) for 4 min per network at 4 Hz with 200 ms exposure of both random and patterned neurons. Calcium videos were processed in ImageJ with manual segmentation and mean fluorescent intensity of each cell was extracted for further processing in MATLAB 2024b. Mean fluorescent intensity traces were pre-processed by removal of the lower envelope with a peak detection bias of 60 s. The traces were then normalized to their baseline (F_0_) using the bottom 10th percentile:

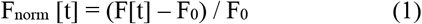

Calcium transient events were detected using findpeaks with a minimum peak width of 500 ms and a minimum peak prominence of 2 standard deviations of the signal and a required 5% ΔF/F magnitude.

Ensemble dynamics were extracted through event coactivity [14]. Significant coactivity was identified as a population coactivity (500 ms bin) greater than a shuffled threshold (shuffle: 1000 permutations of neuron independent randomized temporal-shifting). Significant coactive events (CE) were then identified as ensemble events and were further classified by similarity. We employed a cosine similarity (S):

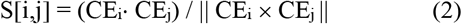

of the neuron feature vectors (e.g. CE_i_ is a vector of 1xN with binary neuron participation of all neurons within the coactive event i) across each ensemble event to determine a likelihood of coactive events occurring from the same population. The map of coactivity similarities was then parsed through principal component analysis, where we identified the number of eigenvalues required to explain over half of the variance as the number of ensembles. Using the number of ensembles, we performed hierarchical clustering on the inverse (dissimilar) map of coactivity similarities with ward linkage. Then clustering of the hierarchical tree was performed by grouping least dissimilar coactive events until the number of ensembles was reached. Ensemble processing scripts are available at https://github.com/Connor-Beck/Dissociated-Ensembles.

### E. Statistics

All data comparisons were tested for normality with the Shapiro–Wilk test. Fabrication and Live-dead datasets were found to be non-normally distributed (p<0.05) and were thus compared using Kruskal-Wallis ANOVA and Wilcoxin Signed Ranks tests respectively. No correction factors were used for either, and the ANOVA test was compared pairwise with a post-hoc Dunn’s test. Mean event rate was computed as the mean number of events over the population of active cells.

## III. Results

### A. Fabrication of meso-circuits

To engineer meso-structured neuronal circuits, we designed patterned substrates featuring localized circular microwells with 150 μm radii, interconnected by 30 μm-wide microchannels to establish global meso-scale connectivity (Figure 1a). We opted for a channel width following previous results of neuronal growth in embossed agarose [12]. Our objective was to examine how neurons self-organize and form connections at both local (intra-microwell) and meso (inter-microwell) levels, providing insights into how architectural constraints influence the emergence of functional connectivity in neural circuits. The 5 microwell configuration offers balance, incorporating localized connectivity within a meso-framework that introduces levels of circuit complexity. Each pattern is confined within a 400 × 400 μm footprint, enabling the implementation of various connectivity paradigms across an individual stamp.

**Figure 1.**
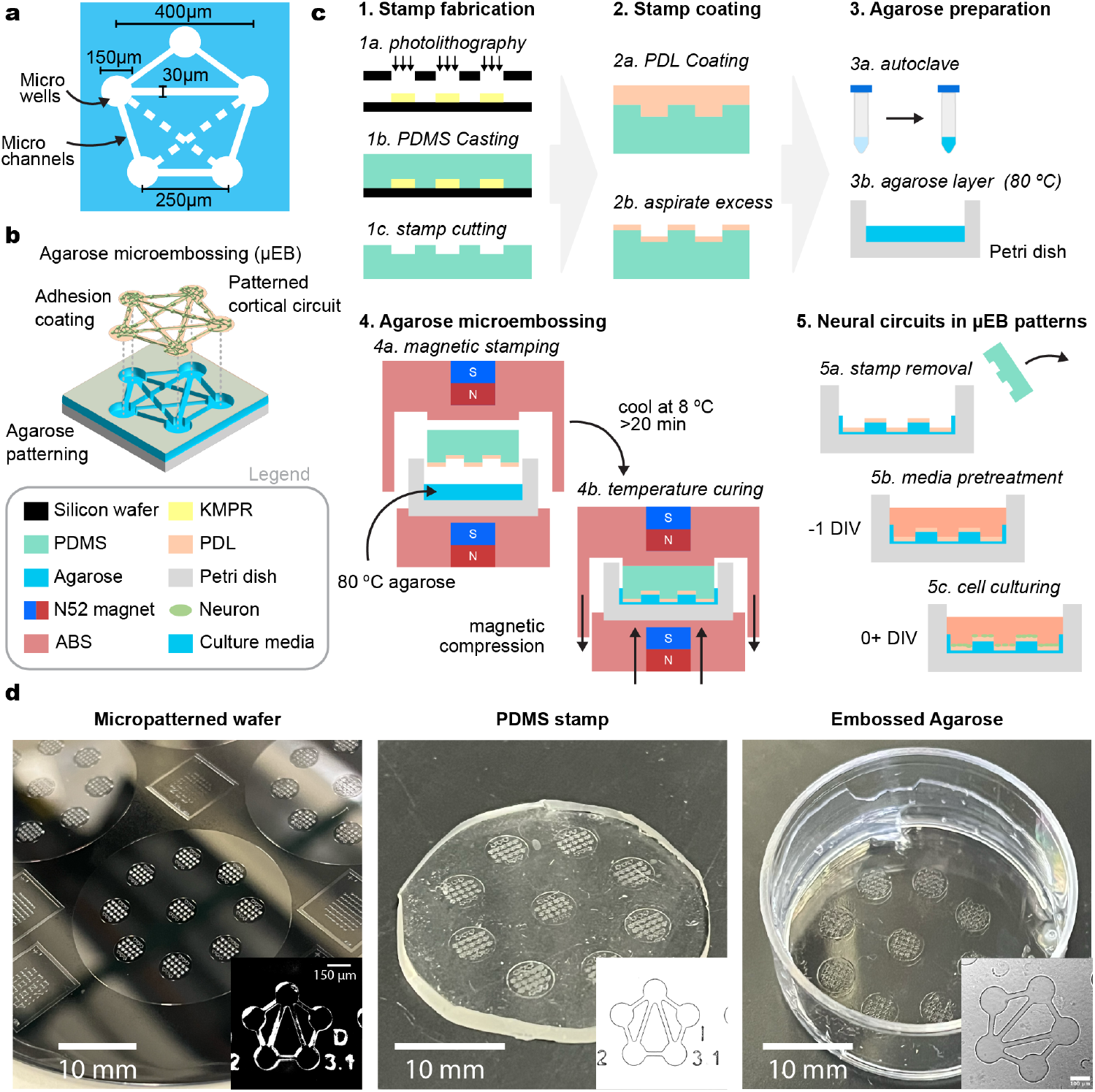
A system for agarose microembossing to introduce patterns for dissociated neural circuits. (a) Schematic design of a single meso-circuit which contains microwells connected through microchannels, that vary across meso-circuits on the chip. (b) 3-D representation of the patterning approach to encourage neural growth and adhesion through hydrogel embossing. (c) Process flow for patterning meso-circuits. (d) Macro and microscopic images across fabrication steps.

To pattern the features, we utilized agarose micro-embossing following previous methods [12]. Here, our goal was to constrain the neuronal growth, promoting network structuring following our patterning design (Figure 1b). Agarose hydrogels enable soft structures for biocompatibility, which can be tuned to mitigate neuronal outgrowth [15], [16]. We selected agarose VIIa at 1% concentration to create the biocompatible devices with rigidity capable of mitigating neurite growth into the agarose. Lower concentrations exhibited reduced pattern fidelity during testing. To perform microembossing, we used soft-microstructures from poly-dimethyl-siloxane (PDMS) inverted patterns structured from a silicon wafer (Figure 1c). To introduce further consistency in batch-to-batch variability, we custom-designed a 3D printable magnetic stamping tool, which harnesses forces from neodymium magnets to create a consistent compressive force.

Altogether, we validated feature formation across the fabrication steps and observed consistent formation of structural features (Figure 1d).

### B. Hydrogel characterization

During embossing, malformation of agarose features was observed (Figure 2a), so we aimed to identify key parameters to produce consistent features. Leveraging the flexibility of magnetic stamping, we sought to optimize the viability of agarose microfeatures as a function of cooling time, thus changing the magnitude of agarose gelation prior to stamp removal. We observed that stamps cooled to 8 °C for ≤14 minutes failed to produce viable features, while a marked improvement in feature fidelity was achieved at 26 minutes of cooling (Figure 2b). Notably, we observed no change in viability between stamps cooled between 20-32 min. A representative non-viable pattern—characterized by incomplete, irregular line formation—is also shown in Figure 2a, in contrast to the sharp and consistent structures of viable designs.

**Figure 2.**
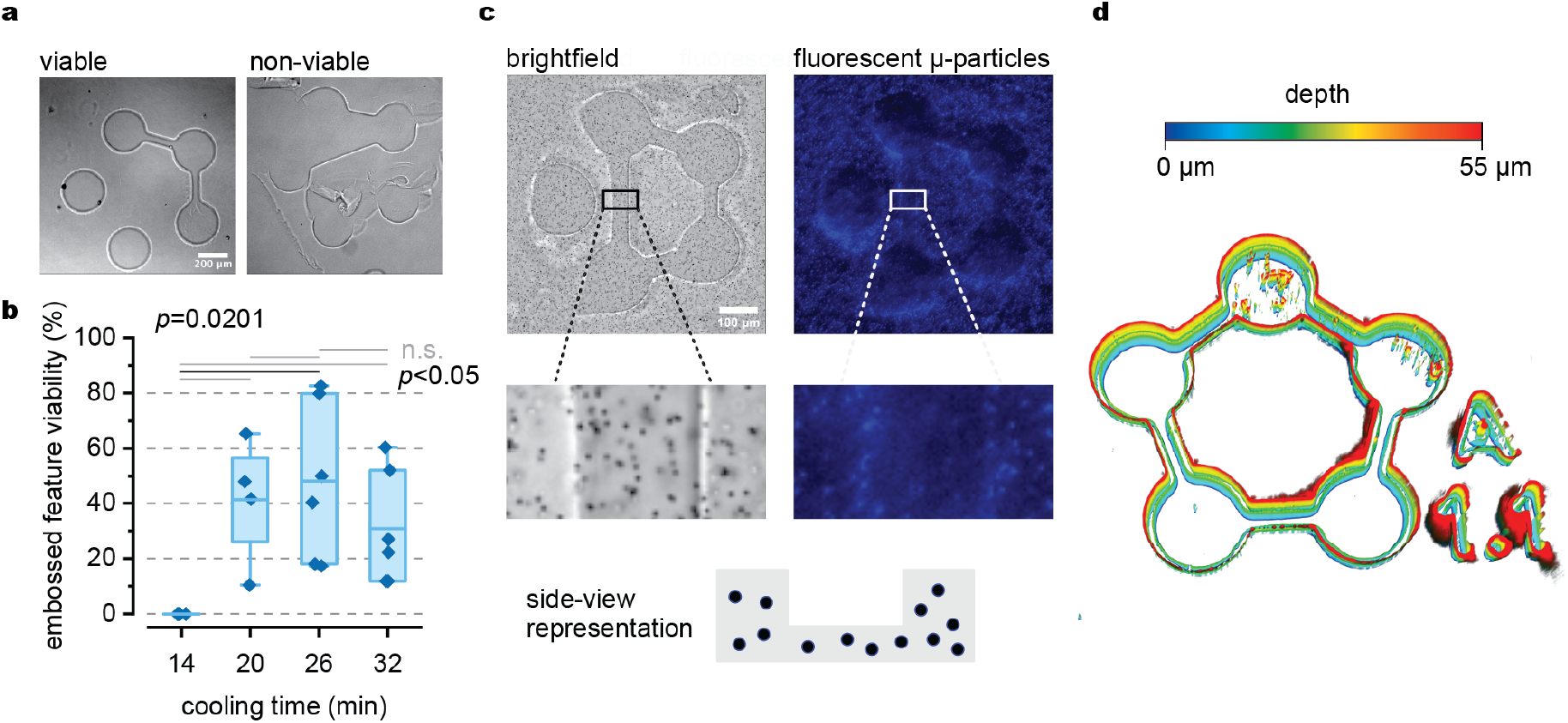
Optimized steps in the method of hydrogel agarose embossing. (a) Microcircuits were visually characterized as viable and non-viable based completeness of features. (b) Different amounts of stamp cooling time led to viable and non-viable features. There was a positive correlation between increased stamp cooling time and feature viability. Statistics are Kruskal-Wallis ANOVA with post-hoc Dunn’s test (n=5). (c) Fluorescent microparticle embedded agarose highlights the 3-dimensional structure of the embossed features, including compressed agarose underneath the features. (d) Contrast adjusted representation of volumetric imaging (Differential interference contrast, z-step: 0.2 µm) shows the measured height of the embossed features.

Following, we tested whether agarose as removed from the bottom of the embossed features, exposing the substrate, or if it was compressed, covering the substrate. To promote neuronal adhesion, an adhesion molecule must be present at the surface, and such variables could impede the use of the device. To investigate this, we incorporated fluorescent microparticles into the agarose, we visualized that agarose settled both in the embossed patterns and the surrounding bulk region. Notably, residual compressed agarose remained on the base of the microfeatures, which could hinder cell adhesion. To address this, we applied poly-D-lysine (PDL) dropwise onto the hydrogel, allowed it to incubate for 5 minutes, and removed excess PDL prior to stamping. This protocol was adapted from microcontact printing techniques known to enhance cell adhesion and proliferation. Neuronal cultures did not adhere well to agarose alone. The PDL coating facilitated neuronal growth by promoting adhesion on the hydrogel surface without compromising biocompatibility (Figure 2c). To confirm the structural integrity of the agarose features, we employed volumetric imaging, revealing consistent feature heights of approximately 50 μm (Figure 2d).

### C. Cell viability in patterned neurons

To characterize the biocompatibility of the microembossed agarose, we used neurons. We cultured primary cortical neurons (E18) to 8 DIV and performed live/dead staining with calcium AM and BOBO-3 iodide fluorescent markers. We counted the number of live and dead cells in individual samples and classified the location of the cells as either in the embossed wells or on the upper surface of the agarose (Figure 3a). There was a significantly greater number of dead cells, but we observed no significant difference between cells situated in the well or on the surface (Figure 3b-c), suggesting that the wells did not reduce cell viability relative to the unconstrained surface.

**Figure 3.**
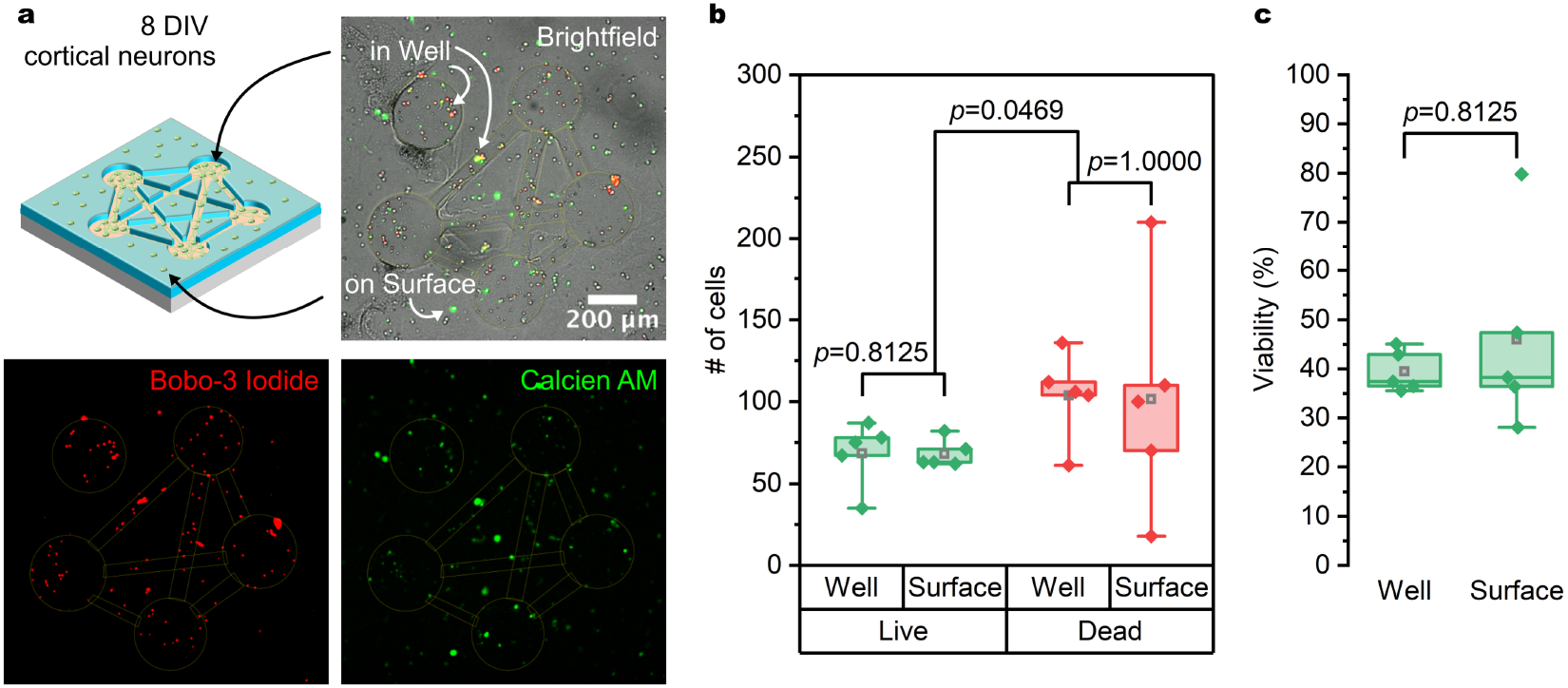
Neuronal cell viability is consistent within the agarose microwells and agarose surface. (a) Representative brightfield and false color fluorescent images (Live: Calcien AM, Dead: Bobo-3 Iodide) of live-dead staining presenting viability results. Same FOV for all three images. (b) Quantification of cells identified as live (soma presenting with Calcien AM and not Bobo-3 Iodide) or dead (Any soma presenting with Bobo-3 Iodide signal). Viability was measured in cells contained within the wells and on the surface (n=5 networks across 4 dishes). (c) Relative cell viability from the data presented in b (Live cell count normalized to the total cell count) shows no significant difference in viability between cells cultured in the well versus the surface of agarose. Data was found to be non-parametric and therefore statistical comparisons were performed with Wilcoxon Signed Ranks test and no correction factors were used.

### D. Neuronal ensembles within patterned circuits

To assess neuronal ensemble formation in our engineered circuits, we performed calcium imaging on dissociated cortical neurons cultured in either random or patterned environments at 14 days *in vitro* (DIV). Recordings over 4 min captured spontaneous activity, allowing us to extract somatic calcium traces across networks (Figure 4a-b). In patterned networks, we observed structured and synchronized activity, in contrast to more dispersed signaling in randomly seeded networks (Figure 4c).

**Figure 4.**
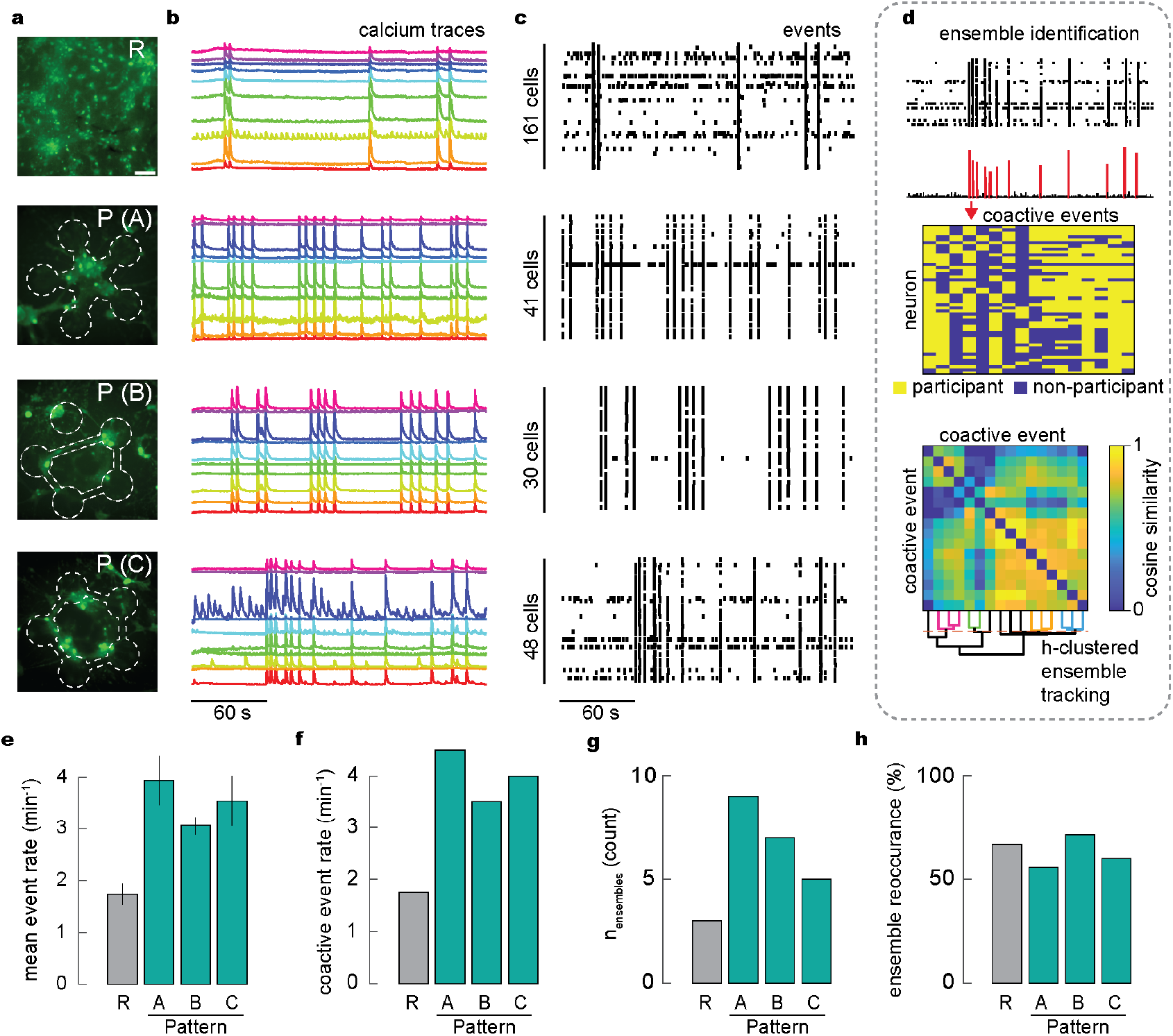
Calcium imaging of patterned meso-circuits reveal formation of distinct ensemble patterns. (a) Representative calcium images (E18, cortical neurons, 14 DIV, Fluo-4AM) of random (R) and patterned (P) networks with differing connectivity patterns (A, B, C). Scale bar: 100 µm. (b) Representative (n=10) mean somatic intensity traces over 4 min of recording. (c) Event raster plots of all active cells identified within each network. (d) Methodological approach for ensemble identification. (i) Ensembles are identified from the population of significant coactive events (>99% shuffled). (ii) Neuron participants between each coactive event are used to map the cosine similarity of coactive events. (iii) Ensembles are identified from hierarchical dissimilarity clustering of cosine similarity matrices. Number of ensembles was identified as the number of eigenvalues required to explain 50% of the cosine similarity variance. (e-h) Measures of (e) neural calcium event activity (mean±SE), (f) network coactivity, (g) number of identified ensembles, and (h) probability of ensemble reoccurrence, measured as the number of ensembles that occur more than once over the 4 min recording window relative to the total number of ensembles.

To detect neuronal ensembles, we implemented a coactivity-based algorithm, identifying coactive events by comparing to shuffled datasets, then implementing cosine similarity to identify repeated coactivity patterns, and hierarchical clustering to localize coactivity patterns into ensembles (Figure 4d). Patterned circuits exhibited more calcium events per cell and greater network-wide coactivity (Figure 4e-f) than random networks, indicating enhanced synchrony. Patterned networks also contained more ensembles (Figure 4g), suggesting that a mesoscale architecture supports functional compartmentalization. Finally, we probed the likelihood of ensemble reoccurrence, defined as the frequency of recurrence of individual ensemble patterns throughout the recording. Ensemble reoccurrence rates were similar across conditions (Figure 4h), indicating stability was unaffected, while patterned networks enabled greater complexity. Overall, patterned connectivity promoted structured, stable neuronal ensembles, supporting the role of physical architecture in shaping functional dynamics.

## IV. CONCLUSION

This preliminary work establishes a robust method for structuring dissociated neuronal networks through agarose microembossing to impose multiscale (micro- and meso-level) connectivity. We demonstrate that these physical architectures not only preserve neuronal viability but also influence network-level dynamics, fostering increased ensemble diversity and synchronous activity. Our approach presents a scalable platform to interrogate how structural constraints shape functional outcomes, enabling studies into neural circuitry without the constraints imposed during in vivo development. Future studies can employ our methodology to study the development of multiscale connectivity or information processing by probing circuit dynamics under external stimuli. This platform holds promise for probing circuit-level phenomena in neurodevelopment, neuropharmacology, and personalized medicine applications.

