## Supplementary Data for "Microembossing Hydrogel Meso-Circuits for Patterning Dissociated Neurons Promotes Ensemble Formation*"

12TH ANNUAL IEEE Engineering Medicine and Biology Society
International Conference on Neural Engineering
11-14 NOV 2025
SAN DIEGO
IEEE EMBS
IEEE Engineering Medicine & Biology Society

### What are neural circuits-on-chip?

**Neural circuit**

- Groups of functionally connected neurons

**Neural circuit-on-chip**

- Neurons grown in a petri dish, which gives a way to analyze the brain's function outside of the body

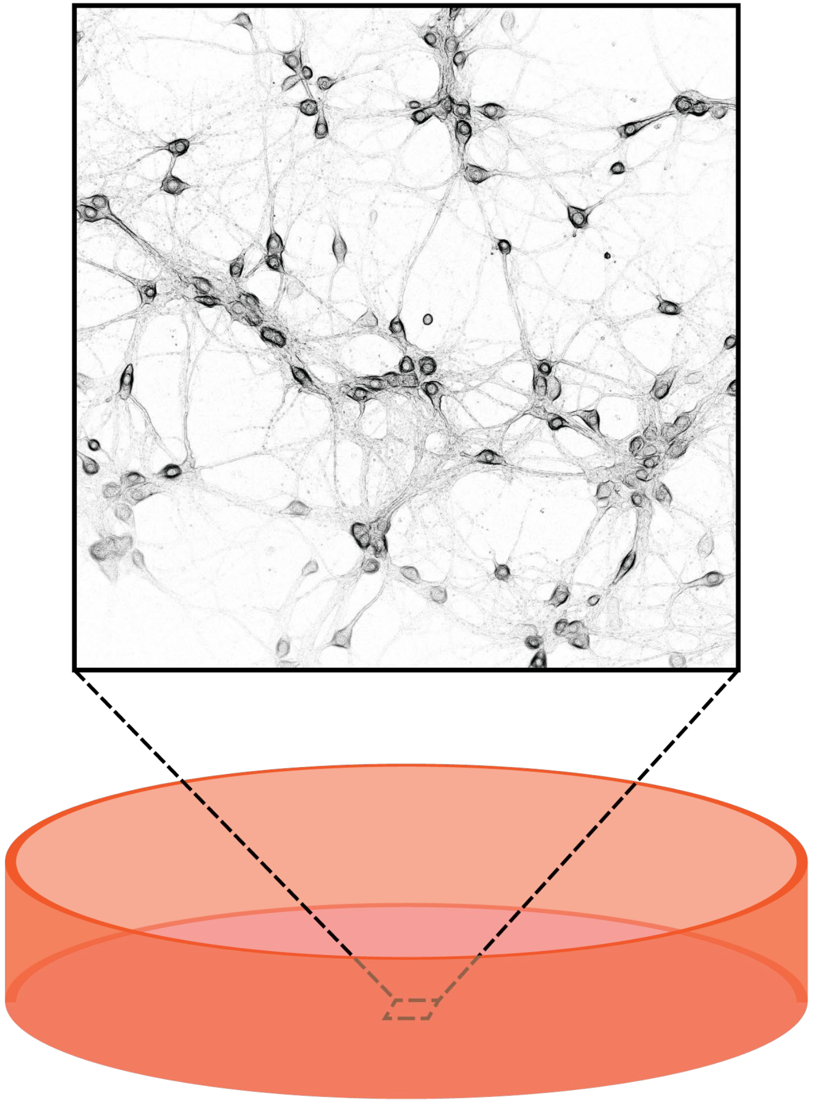

### Why engineer neural circuits-on-chip?

**Potential for early-stage pharmaceuticals and personalized medicine**

- Low cost
- Fast timelines
- Engineered control
- Experimental access

**However, you lose coordinated functions representative of *in vivo***

**Hypothesis:** Patterning dissociated cortical circuits restores brain-like dynamics.

[Chow Neurochem. 2022]

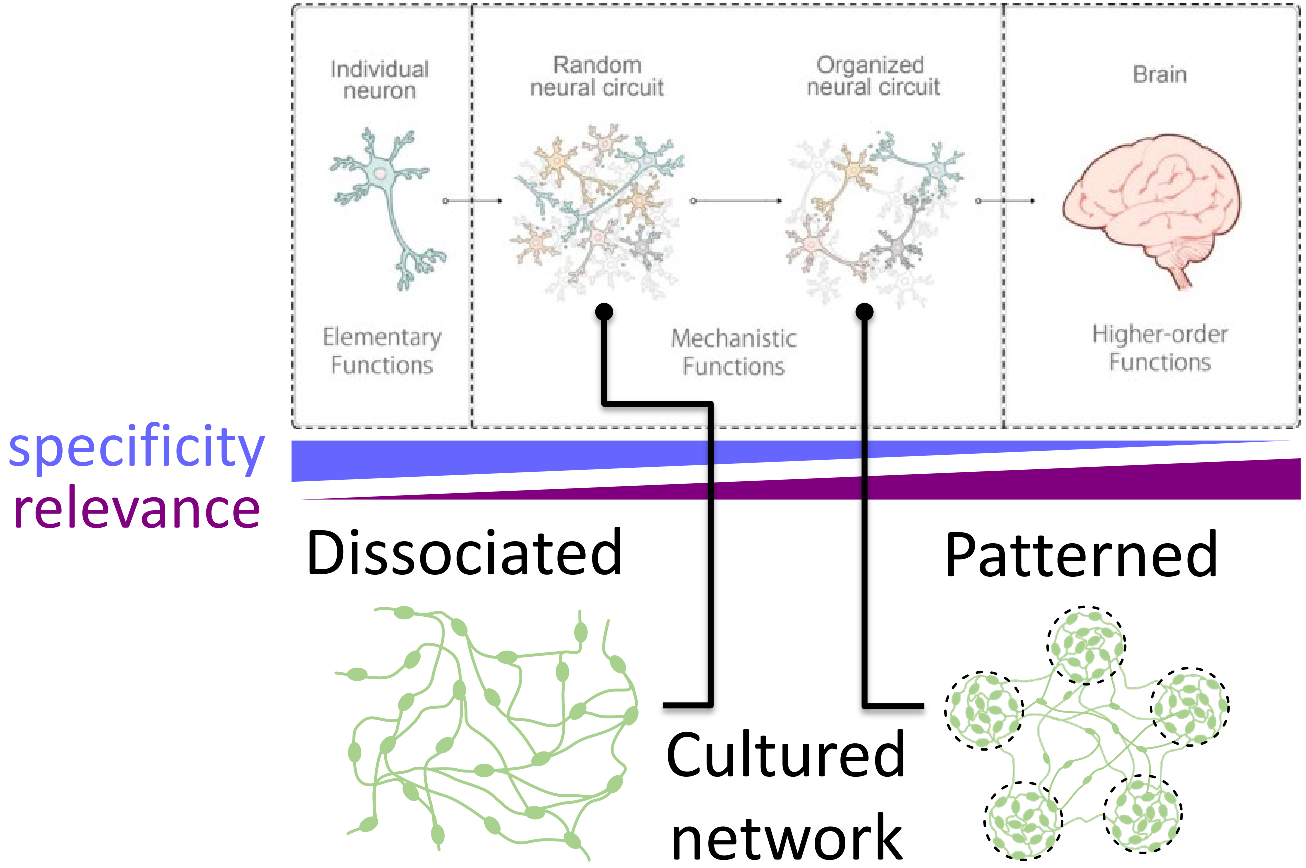

### What is missing in neural circuit-on-chip?

**Observation 1.**
Coordinated functions arise as cortical circuits connect across scales

**Observation 2.**
A crucial identity of coordinated function in the cortex is *ensemble firing*

**Observation 3.**
Ensembles are lost *in vitro*

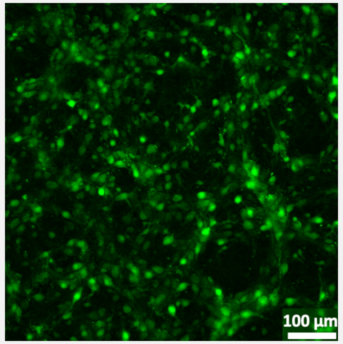


Calcium events of a randomized cortical network

coactive events are non-unique population wide events

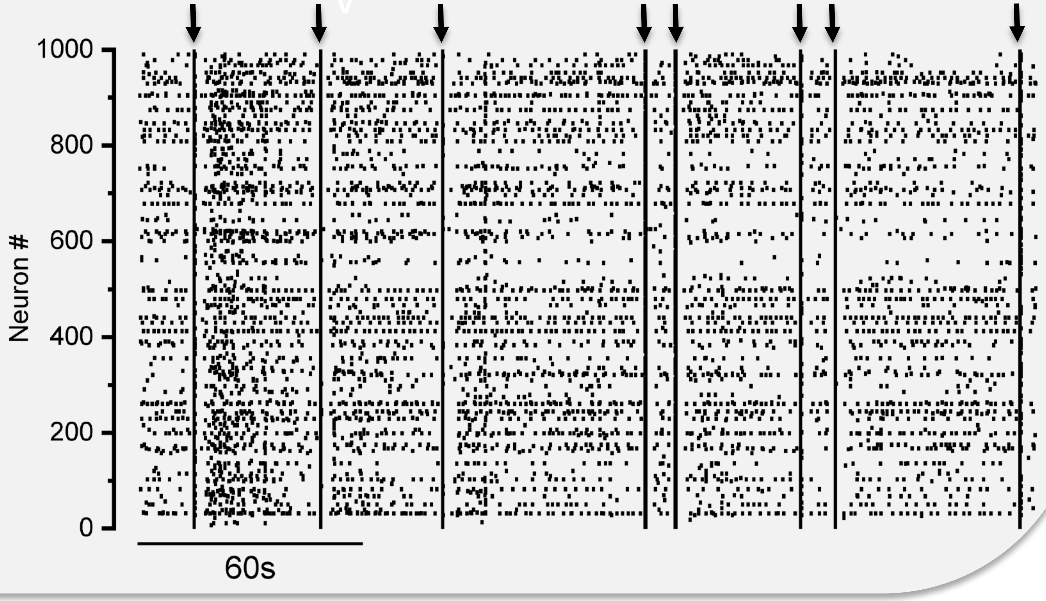


*Ensembles are groups of neurons firing in coordinated patterns*

**To bridge the gap from cell cultures to *in vivo***, introducing circuit complexity is essential.

Engineering multi-scaled connectivity

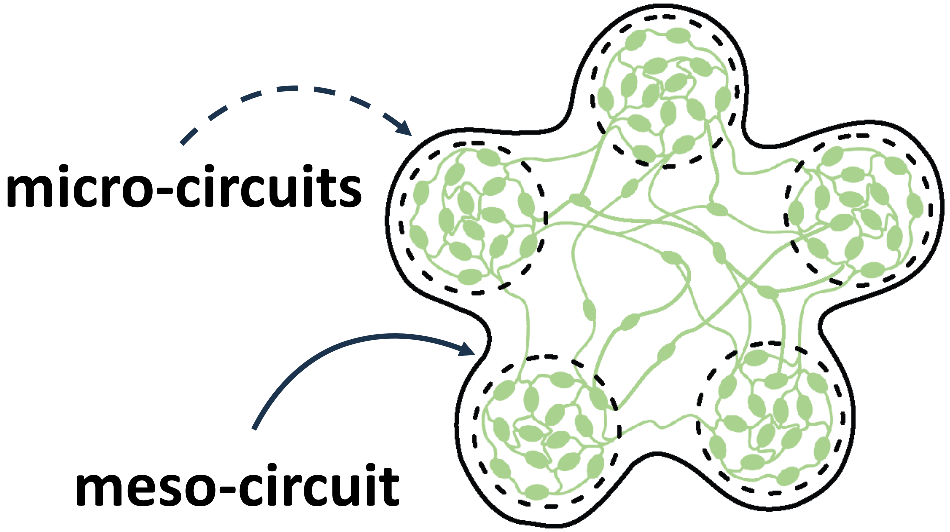

### Creating an easy-to-use device

We need to engineer **robust, reproducible,** and **easy-to-use** methods to enable brain-like circuits *in vitro*

Engineering development

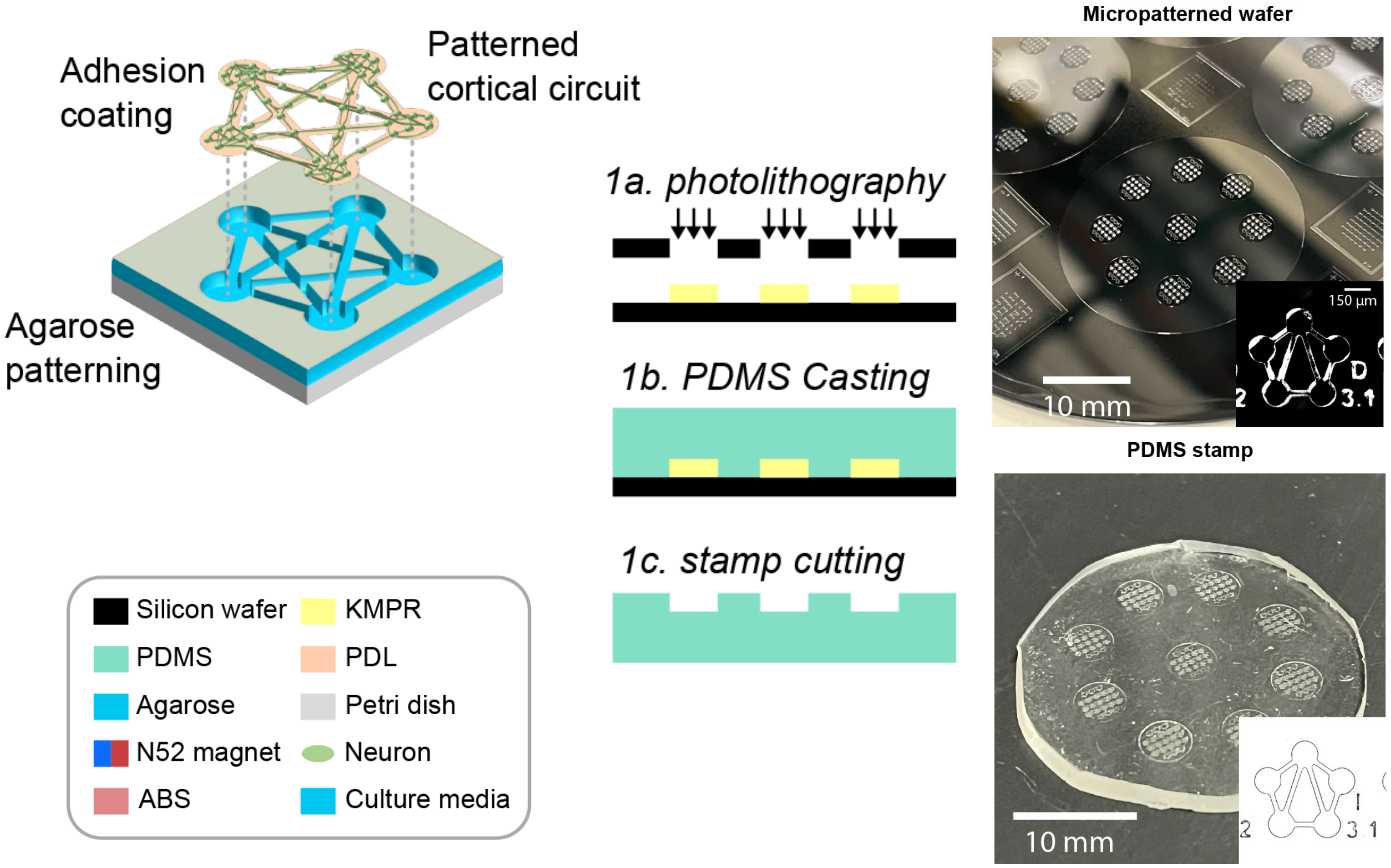


#### Reproducible neural circuit patterning with hydrogels

1. Cleanroom fabrication
2. PDMS stamp forming
3. Hydrogel embossing

*Let's make accessible to all cell-culture facilities*

### Easy-to-use agarose hydrogel embossing

**1.** Prepare PDMS and agarose.

**2.** Use magnetic stamping tool to emboss patterns during gelation.

**3.** Lift stamp and prepare patterned agarose for cell seeding.

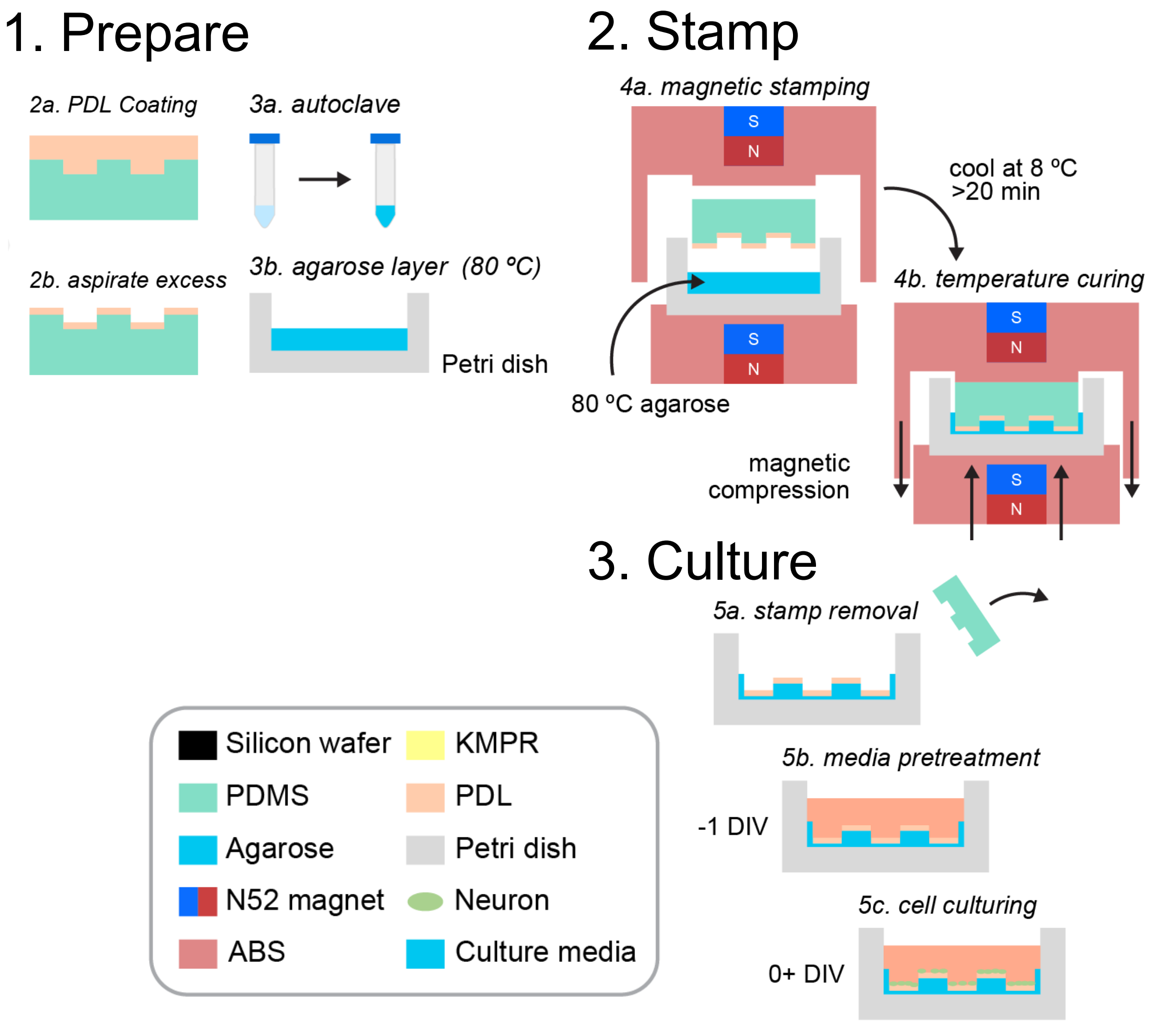


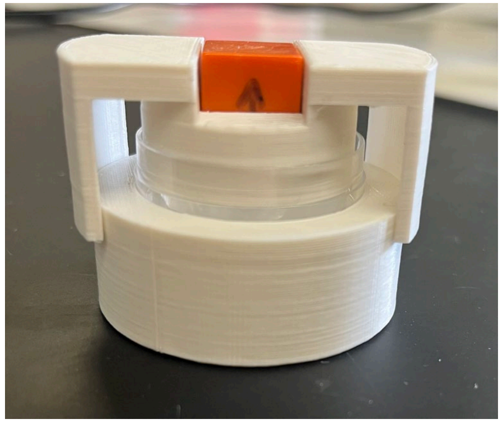

Embossed Agarose

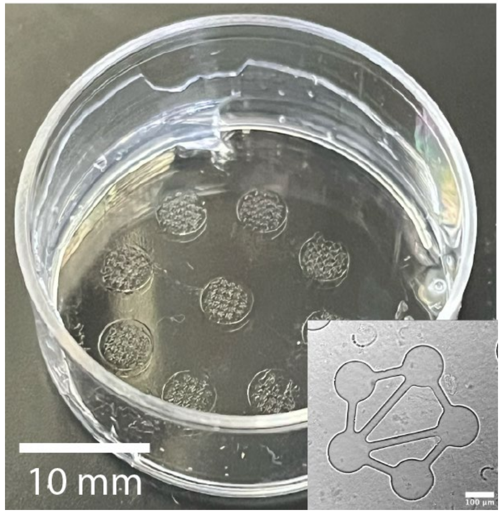

### Establishing methodologies for usability

1. Brief cooling caused incomplete or collapsed pattern formation.
2. Fluorescent microparticles embedded in the agarose show thin layer within embossed patterns

#### Engineering reproducibility

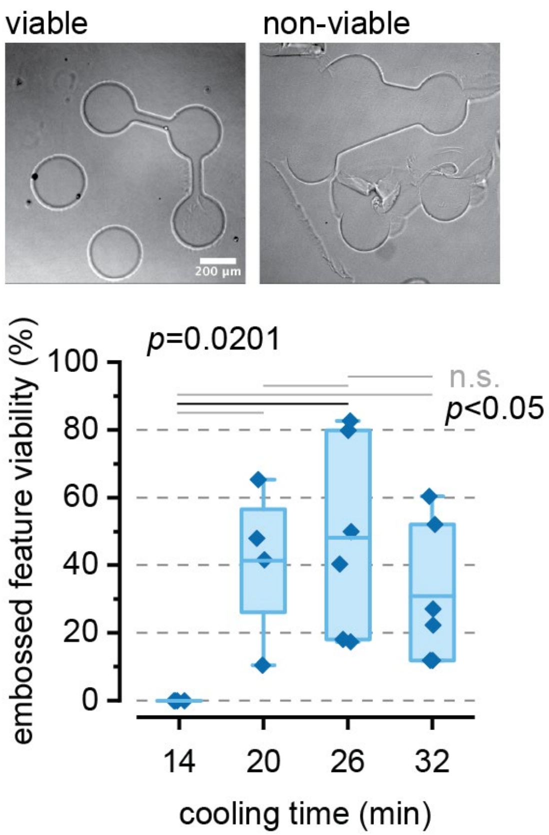


#### Agarose resides across the interface

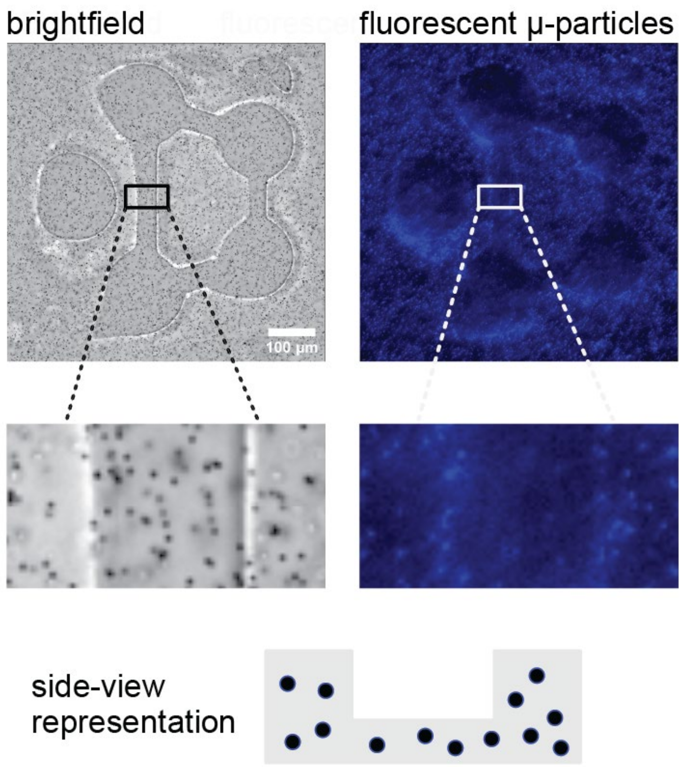

### Neural viability on agarose

We found no significant difference in cell health found between neurons within the patterned features and the surface (Mann-Whitney U test)

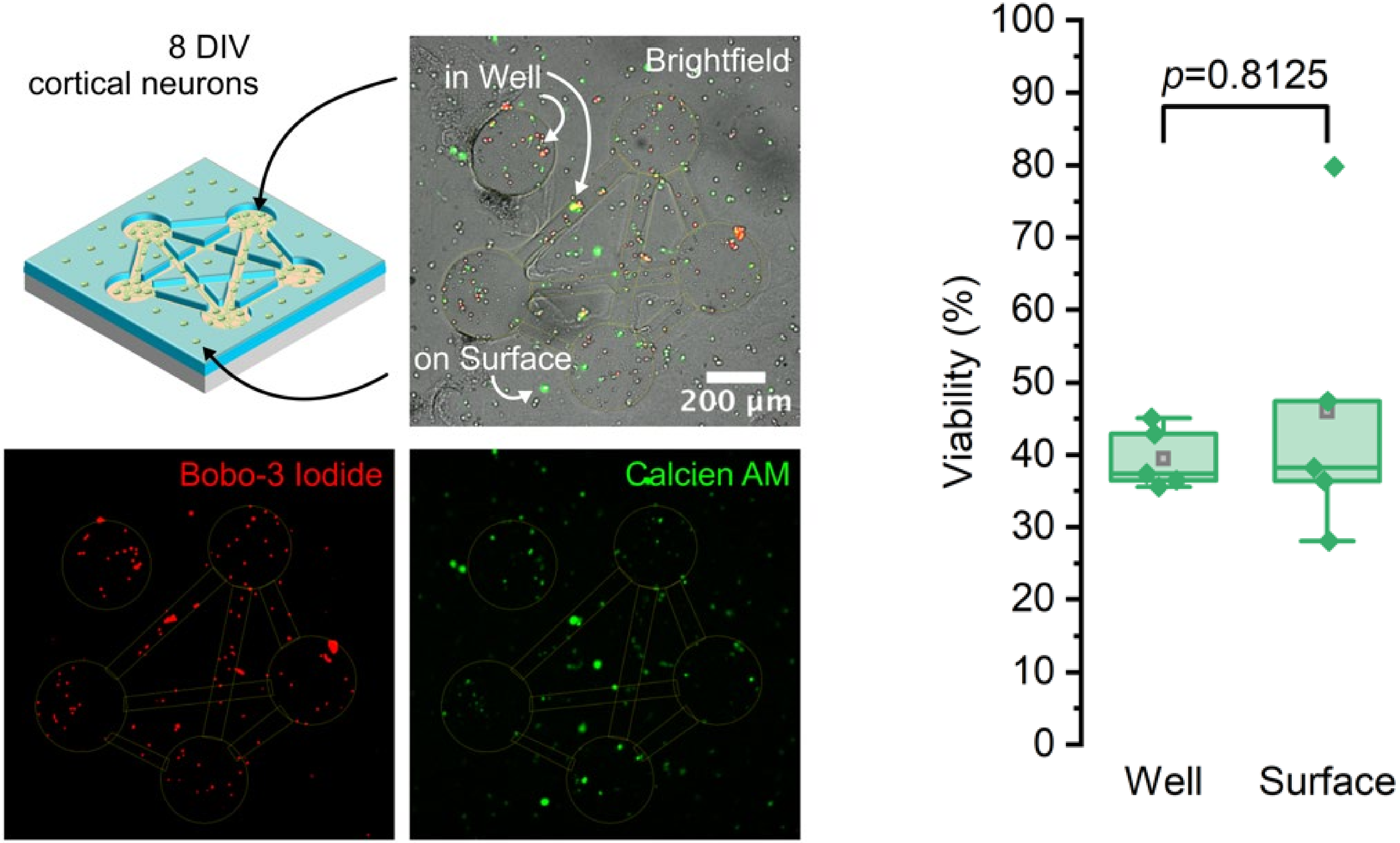


**Primary E18 rat cortical neurons cultured to 8 DIV in developing circuits**

### Neuronal Calcium  Dynamics in Agarose Embossed Features

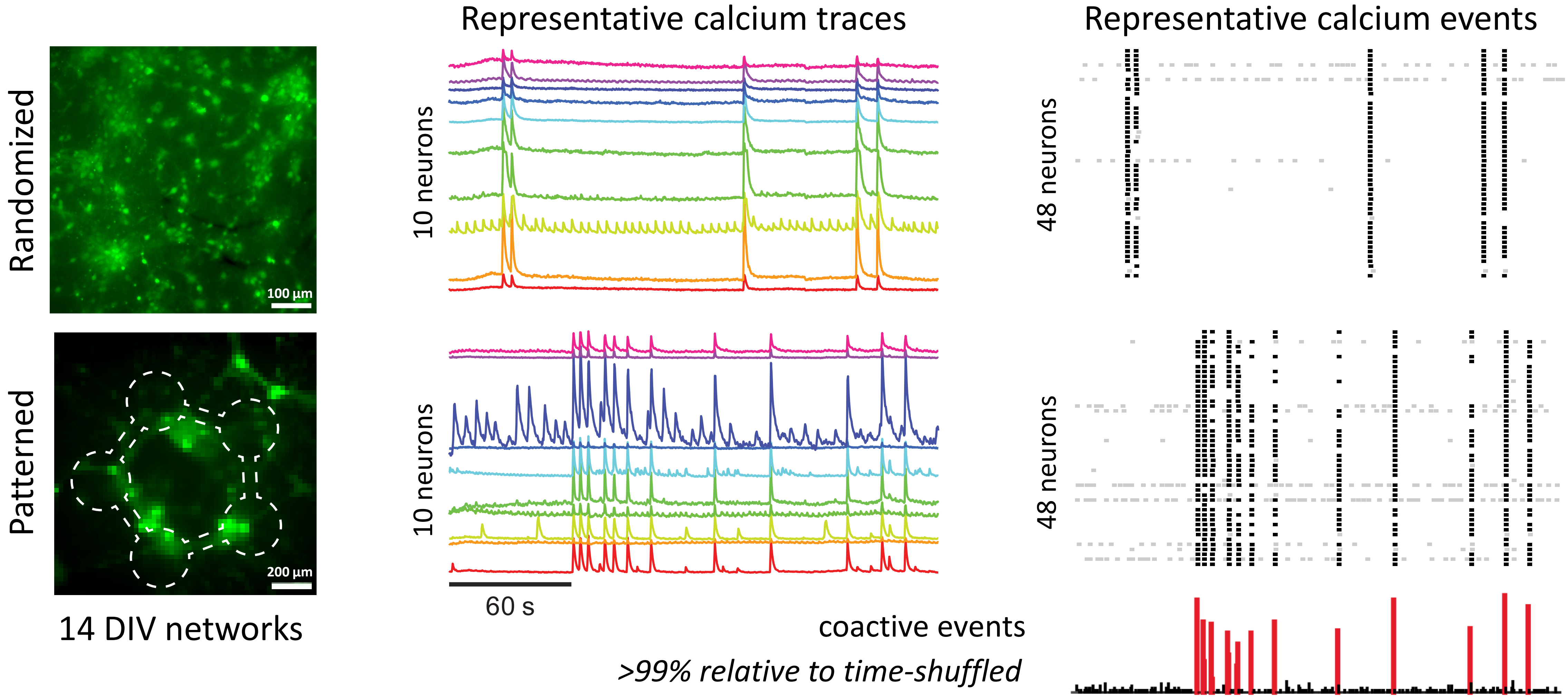

### Coactive network features seen across patterned networks

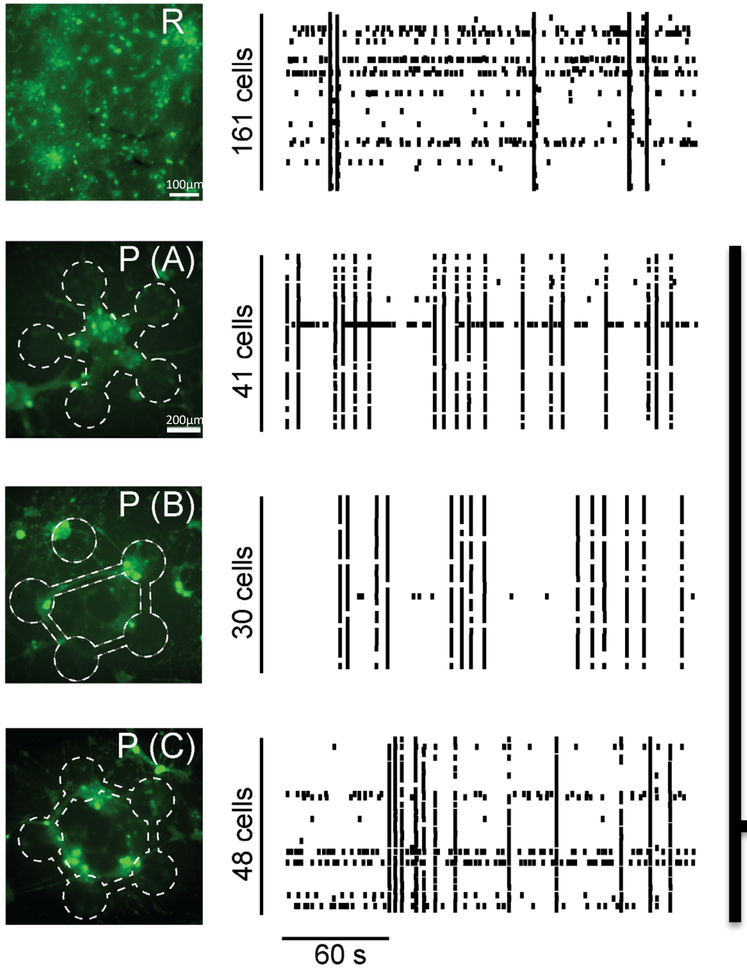


single neuron values

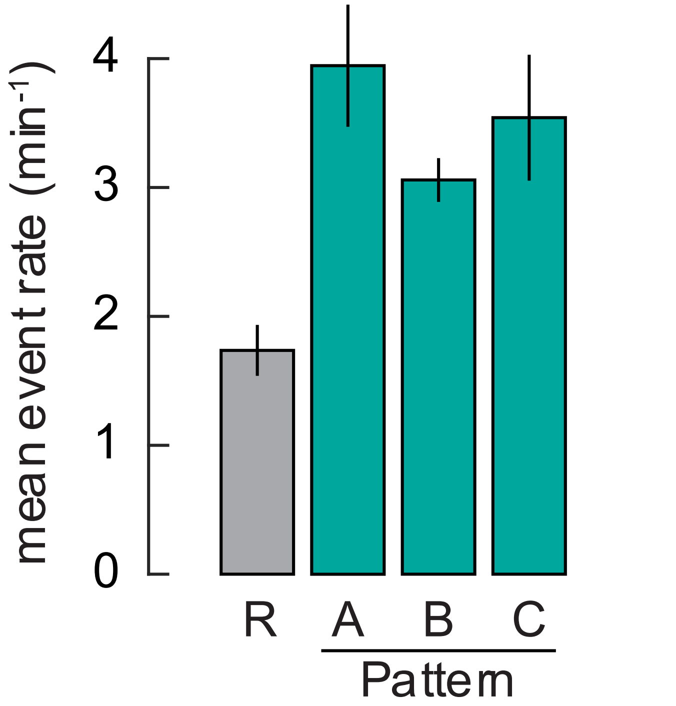


network value

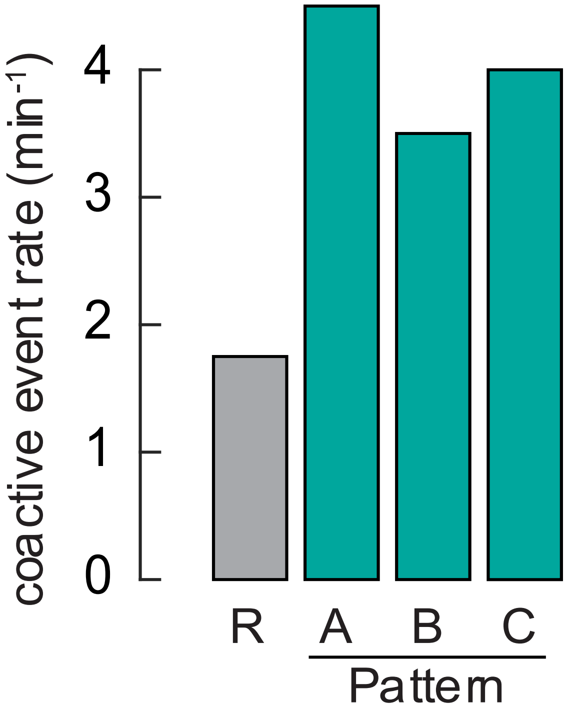


Are these coactive events unique codes?

### Unique ensemble codes arise in patterned networks

#### Identifying ensembles from coactive events

For more information on ensemble identification see [Wenzel and Hamm, J Neuro methods 2020]

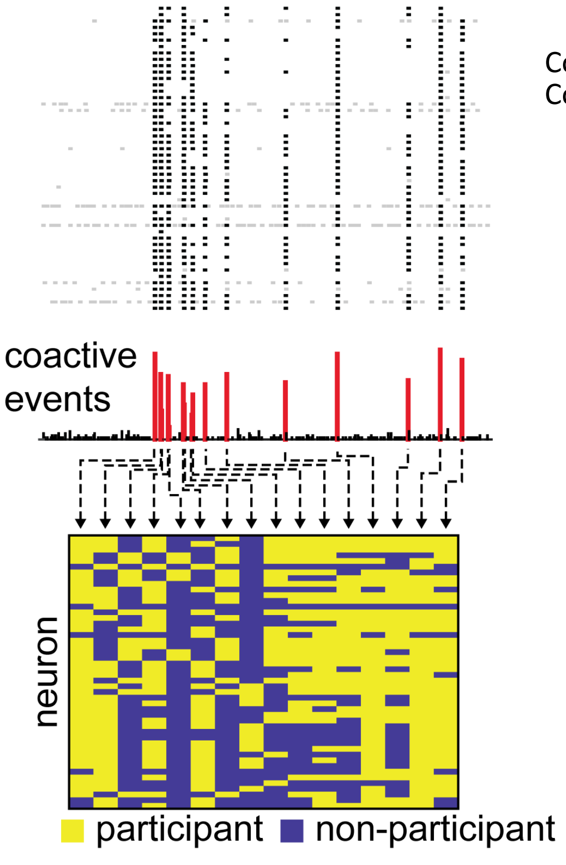


“neural barcode”

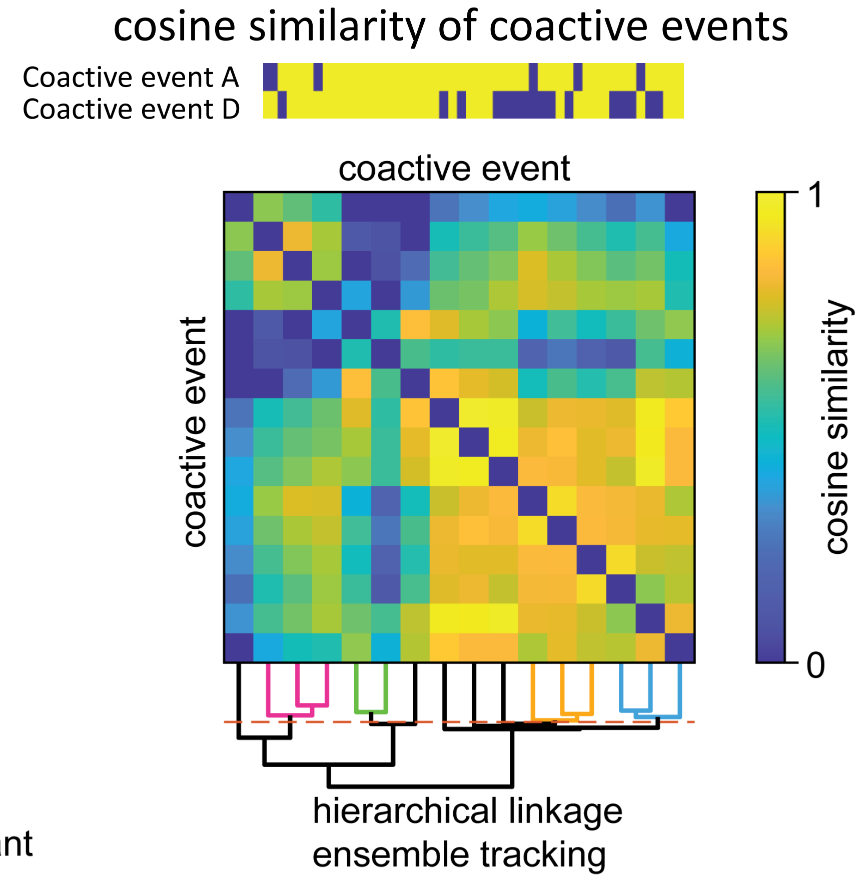


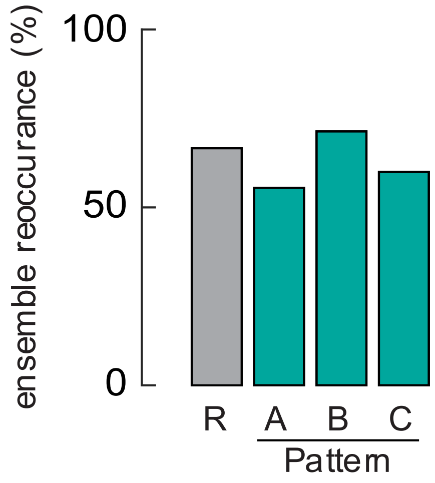


**Patterned ensembles repeat similarly to random networks**

**More ensembles in patterned networks**

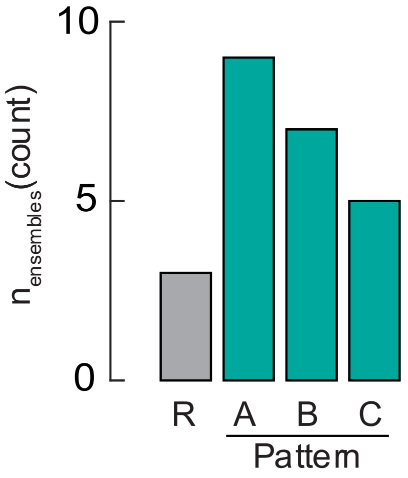

### Conclusions

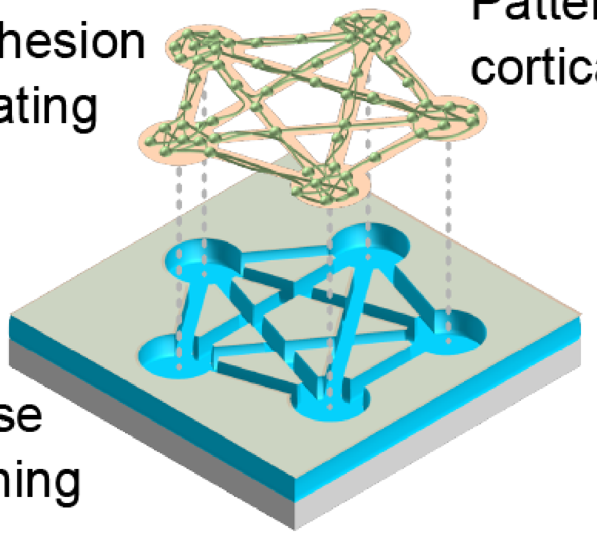


- Developed a **robust, reproducible, and easy-to-use** agarose embossing workflow for neuronal circuits.
- Demonstrated **multiscale architecture and rich ensemble dynamics**.
- Provides a new platform to study:
  - Multiscale circuit formation
  - Applications in pharmacology and personalized medicine

### Thank you!

#### Microembossing Hydrogel Meso-Circuits for Patterning Dissociated Neurons Promotes Ensemble Formation

Mckennah Thompson, Connor Beck, and Anja Kunze; IEEE NER 2025

**Thanks to all involved**

This research is supported by the Montana Microfabrication Facility, NSF CAREER AWARD (#CBET-1846271, AK) and Montana State University Undergraduate Scholars Program (MT)

**Mckennah Thompson**
**Dr. Connor L. Beck**
**& Dr. Anja Kunze**

Montana State University
Kunze Neuroengineering Laboratory

Contact:

Research: https://www.kunzelab.org
Follow us on X: @TheKunzeLab
